# Neural correlates differ between crystallized and fluid intelligence in adolescents

**DOI:** 10.1101/2024.10.06.616909

**Authors:** Bowen Qiu, Rui Qian, Baorong Gu, Zhifan Chen, Zichao Li, Mingyang Li, Dan Wu

## Abstract

Fluid and crystallized intelligence are acknowledged as distinct facets of cognitive ability during brain development, but the specific neural substrates and molecular mechanisms underlying them remain unclear. This study used a sample comprising 7471 young adolescents (mean age 9.87 ± 0.62 years) from the ABCD cohort to elucidate the differential neural correlates of fluid and crystallized intelligence. Our findings indicated that micro-level brain MRI phenotypes such as water diffusivity were closely associated with fluid intelligence, whereas macro-level brain MRI phenotypes such as gray matter cortical thickness were indicative of crystallized intelligence. We further investigated the molecular mechanisms underlying fluid and crystallized intelligence by correlating the characteristic MRI markers with spatial transcriptome profiles and PET imaging. Results showed that fluid intelligence had significant associations with serotonin and glutamate system, while crystallized intelligence was related to serotonin, dopamine and acetylcholine system. Furthermore, we examined the impacts of lifestyle factors on these two forms of intelligence and how the molecular pathways mediated these impacts. Our investigation suggested that physical activities, screen use and sleep duration influenced fluid intelligence mainly through mGlu5 receptors and crystallized intelligence through 5HT1a and D2 receptors. In conclusion, these findings illustrated a distinct neural basis between fluid and crystallized intelligence from the perspectives of neuroimaging, neurotransmitters, and lifestyles in young adolescents.

## Introduction

Comprehending the mechanism of general intelligence has been an important area of research for over a century ^1^. General intelligence can be decomposed into fluid and crystallized intelligence ^2,3^. Fluid intelligence is characterized by the capacity to reason and solve problems in novel situations, reflecting the ability to adapt and learn independently of prior knowledge. In contrast, crystallized intelligence is shaped by accumulated experiences and learning, representing knowledge and expertise developed over time ^4^. Distinguishing between fluid and crystallized intelligence is vital for understanding how individuals approach cognitive tasks, highlighting the dynamic interplay between adaptability and accumulated knowledge in human cognition. In adolescents, the development of fluid and crystallized intelligence follows distinct trajectories and fundamentally shaped brain development ^5,6^.

Many studies tried to identify differences between two forms of intelligence using neuroimaging technology ^7,8^ and identified several imaging characteristics ^9,10^. Fluid intelligence was found related to the executive network within the dorsolateral prefrontal cortex, anterior cingulate gyrus, and inferior and superior parietal lobules ^11,12^. Higher crystallized intelligence level was found associated with cortical volume and thickness of the whole brain ^13–15^. Moreover, fluid and crystallized intelligence both had significant associations with white-matter tract integrity and microstructural properties, which might be related to information processing ^16–18^. However, these studies mainly focused on adults ^19^, and thus could not intuitively reflect the basis of adolescent intelligence.

Several studies have reported that neurotransmitter systems were associated with general intelligence using tools such as blood biochemical tests, DNA analysis, and positron emission tomography (PET). The neurotransmitter systems included serotonin system ^20–22^, dopamine ^23–25^, acetylcholine ^26,27^, GABA ^28,29^ and glutamate ^30,28^. Only a limited number of studies have attempted to examine a specific form of intelligence ^21,31^. Due to the lack of separate quantification of fluid and crystallized intelligence, previous research on molecular mechanisms did not distinguish the neurotransmitter basis of fluid and crystallized intelligence ^8^.

Besides the molecular drivers, lifestyle was known to have a significant impact on adolescent intelligence ^32,33^, and 24-Hour Movement Guidelines for Children and Youth were suggested to promote mental health and intelligence development ^34–37^. The guidelines recommended daily physical activity, adequate sleep duration, and limited screen time to promote overall well-being and cognitive development in children and youth. Previous studies indicated that physical activities ^38,39^ had a positive effect on adolescent intelligence. Meanwhile, screen use was traditionally believed to have negative effects ^40,41^, and the causal effect was recently confirmed through the method of Mendelian Randomization ^42^. Short sleep duration was also shown to hurt intelligence development ^43–45^. However, the molecular mechanisms underlying the influence of those lifestyles on intelligence have not been well understood.

In this study, we aimed to differentiate the neuroimaging phenotypes between fluid and crystallized intelligence and to further identify the neural and genetic mechanisms supporting the two forms of intelligence. We used neuroimaging and cognitive data from the Adolescent Brain Cognitive Developmental (ABCD) study. The pipeline of the current study is illustrated in Figure 1. We first evaluated the association between neuroimaging phenotypes and each form of intelligence measure to obtain intelligence-associated MRI maps using the linear mixed model (Figure 1b). We then performed spatial correlation between the MRI-intelligence association maps and Allen Human Brain Atlas (AHBA) transcriptome maps to identify the highly related genes, followed by genetic enrichment analysis (Figure 1c). Furthermore, we correlated the association maps with neurotransmitter density maps from PET (Figure 1d). Finally, mediation analysis was utilized to uncover the association between lifestyles, MRI phenotypes, and intelligence, as well as the underlying neural basis (Figure 1e).

**Figure 1:**
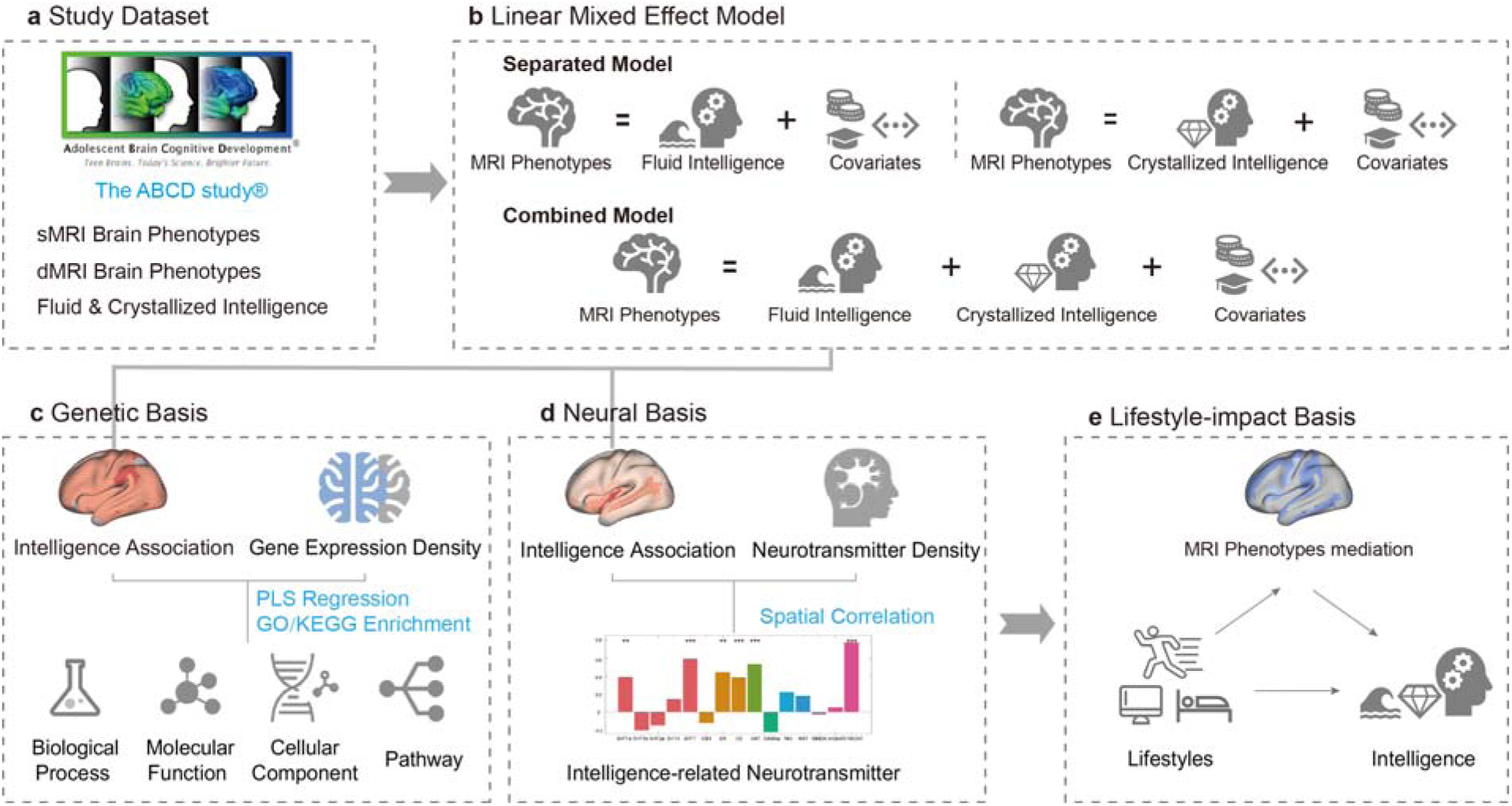
The study workflow. (a) Study datasets and main measures used in the present study; (b) Two different linear mixed models (LME). Separate model investigated the direct relationship between MRI phenotypes and two forms of intelligence, while combined model investigated the relationship between MRI phenotypes and one intelligence after removing the influence of the other one; (c) Spatial correlation between intelligence-associated MRI maps and AHBA transcriptome maps, followed by GO/KEGG enrichment; (d) Spatial correlation between intelligence-association MRI maps and neurotransmitter density maps; (e) Spatial correlation between lifestyle-mediation maps and neurotransmitter density maps.

## Methods

### Participants and datasets

#### The ABCD study

The ABCD study was a longitudinal population study to investigate the development of a diverse group of young adolescents, aged 8-18 years, from various sites across the United States (source: https://abcdstudy.org). Each participating center ensured that they obtained comprehensive written consent from parents and enthusiastic assent from the children involved. Rigorous research protocols were reviewed and approved by the institutional review board of all participation sites. For this investigation, we utilized data from the ABCD Data Release 5.0. The present analysis included a sample of 7045 individuals with complete demographic information, intelligence assessments, and MRI data.

#### Intelligence measures

Crystallized intelligence and fluid intelligence were evaluated using the NIH Toolbox ^46,47^, a widely recognized tool known for its strong reliability in assessing children (Akshoomoff et al., 2013). The toolbox encompassed seven tasks: picture vocabulary, flanker inhibitory control and attention, list sorting working memory, dimensional change card sort, pattern comparison processing speed, picture sequence memory, and oral reading recognition. The NIH Toolbox yielded three composite scores for crystallized, fluid, and total intelligence ^47^. For this study, we used corrected t-scores of the crystallized and fluid intelligence.

#### Neuroimage data

In the ABCD study, 3D T1- and T2-weighted structural images along with diffusion images were obtained using 3T scanners across 21 data collection sites ^48^. The scanning and preprocessing details have been previously outlined ^49^. This study used the preprocessed data including macro-structure such as brain volume (BV), surface area (SA), and cortical thickness (CT); as well as micro-structure such as fractional anisotropy (FA), mean diffusivity (MD), longitudinal diffusivity (LD) and transverse diffusivity (TD) in the Destrieux cortical atlas (Destrieux et al., 2010).

#### Gene expression data

The human microarray-based gene expression data were obtained from the AHBA (http://human.brain-map.org) ^51^. These data were sourced from six postmortem adult brains (1 Hispanic, 2 African-American, and 3 Caucasian; aged 24–57 years, comprising one female and 5 males). Specifically, AHBA provided comprehensive coverage of nearly the entire brain, including normalized expression data of 20,737 genes with unique Entrez IDs detected by 58,692 probes obtained from 3,702 spatially distinct tissue samples.

Given the large transcriptional differences between cortical and subcortical regions, we performed gene expression analysis within the cortical regions. Then we used the abagen toolkit (https://www.github.com/netneurolab/abagen) ^52^ based on Python to extract the transcriptome matrix at the region-of-interest (ROI) level, and ROI partitioning was based on Destrieux2009 atlas ^50^. The processing pipeline was summarized in ^53^. As a result, we retained a total of 15,632 genes for further analysis, resulting in a gene expression matrix of 148 regions × 15,632 genes.

#### Neurochemical data

The neurochemical data were derived from the average group maps of healthy volunteers who had participated in prior PET and SPECT studies, on fifteen receptors and transporters ^54^, spanning eight neurotransmitter systems, including serotonin (5HT1a, 5HT1b, 5HT2a, 5HT4, and 5HTT), dopamine (D1, D2, and DAT), GABA (GABAa), μ-opioid (MU), norepinephrine (NAT), cannabinoid (CB1), acetylcholine (VAChT) and glutamate (mGluR5 and NMDA). These density maps, available from an online repository (https://github.com/juryxy/JuSpace/tree/JuSpace_v1.5/JuSpace_v1.5/) were normalization to the Montreal Neurological Institute (MNI) space. Subsequently, the maps were linearly rescaled to a standardized range of 0–100 ^54^. We calculated the mean value of the neurotransmitter data in the 148 regions based on Destrieux2009 parcellation, and the results were then used to perform the spatial correlation with intelligence-related MRI markers.

#### Lifestyle data

The lifestyle data were provided by the ABCD study ^45,55^. Based on the 24-Hour Movement Guidelines, we included three kinds of lifestyles (physical activity, screen use, and sleep duration) from the ABCD Risk Behavior Survey Exercise Physical Activity, Screen Time Questionnaire (Parent report), and Sleep Duration Scale for Children (Parent report). The value of physical activity represented how many days the teenager was physically active over 60 minutes during the past 7 days. Screen use was measured as the average hours an adolescent used screen in a day. Sleep scores, ranging from 1 (more than 9 hours) to 5 (less than 5 hours) represented the adolescent sleep duration, where higher scores meant shorter sleep duration.

### Statistical analysis

#### Linear mixed effect analysis

The linear associations between fluid/crystallized intelligence and structural MRI (sMRI)/ diffusion MRI (dMRI) phenotypes were examined using a linear mixed-effect model (LME). Considering the high correlation between the two forms of intelligence, examining them separately as reported in previous studies ^10,56^ may not be able to disentangle their intertwined effects. We applied two LME models by modeling the fluid and crystallized intelligence separately or jointly.

In the separate LME model, we included age, sex, ethnicity, race, family income, and parental education as fixed effects, while also factoring in the MRI site as a random effect. In the combined LME model, the other form of intelligence was added to the fixed effects.

Furthermore, brain regions demonstrating significant associations with intelligence (p < 0.05) were utilized to construct intelligence-MRI association maps ^57^, based on the association coefficients in the combined LME model. These maps were then utilized for spatial correlation analysis in subsequent investigations.

#### Partial Least Regression Analysis

PLSR was a widely used multivariate method to identify relationships between multiple predictor and response variables ^58,59^, and it was widely used in gene enrichment ^53,60^. Here we utilized PLSR to explore intelligence-related genes. In the PLSR model, the predictor variables were the z-statistics normalized matrix of the sample-wise mean gene expression level of all probes from each donor. The response variables were the intelligence-MRI association maps.

After the analysis, PLSR components were ranked based on explained variances, and thus, the components with the highest explained variances represent the covariance of high-dimensional data matrices through low-dimensional sequences. We used permutation tests (permutation times = 5,000) to assess the statistical significance of each PLSR component ^61^. For each significant component, we used a bootstrapping method to correct the estimation error of the weight of each gene ^62^. We then ranked the genes according to their corrected weights, which represented their contribution to the PLSR component.

#### GO/KEGG enrichment

Gene Ontology (GO) ^63^ and the Kyoto Encyclopedia of Genes and Genomes (KEGG) ^64^ were employed for functional annotation of the intelligence-related genes, encompassing both positive and negative gene weights. Genes with top 10% positive weight were selected for enrichment ^65,66^.

For gene set functional enrichment analysis, we first utilized the GO and KEGG annotations of genes from the annotation package org.Hs.eg.db (version 3.1.0) to map genes into the background set. Subsequently, we conducted enrichment analysis using the R package clusterProfiler ^67^ to obtain the results of gene set enrichment. All enrichment analyses were performed using Fisher’s exact tests and adjusted p-value < 0.05 together with q-value < 0.05 ^59,62^ were considered statistically significant.

#### Atlas-based spatial correlation

Based on the intelligence-MRI association maps, atlas-based spatial correlations based on Destrieux ROIs were applied through the JuSpace toolbox ^54^. In this study, intelligence-MRI association maps were correlated with PET/SPECT-neurotransmitter density maps to evaluate the neurotransmitter basis underlying the two intelligences. We applied Spearman’s rank correlation and used bootstrapping to estimate the significance. The significance was tested through 5000 permutations ^68,69^. Additionally, precise p-values for spatial correlation were calculated by recomputing the correlation using null t-statistic maps obtained through label shuffling for intelligence levels ^49^.

#### Mediation analysis

We applied mediation analysis using the Matlab Mediation toolbox ^70,71^ to examine whether and how the impact of lifestyle on intelligence was mediated by the MRI phenotypes. For each brain region, the Average Mediation Effect (AME) was used to represent the mediating effect. Similar to the method above, the significant AME map was also spatially correlated with the neurotransmitter density map to elucidate the specific neurotransmitters supporting the mediation effect.

## Results

### Overview of fluid and crystallized intelligence in Adolescents

In the ABCD study, 7471 participants (aged 9.87 ± 0.62 years old, 3538 [47.3%] females) underwent intelligence testing using the NIH Toolbox ^46,47^ and had complete, preprocessed sMRI and dMRI data ^72^. Intelligence measures included fluid intelligence (fully corrected t-scores: 96.25 ± 17.29) and crystallized intelligence (fully corrected t-scores: 107.09 ± 18.43).

Fluid and crystallized intelligence showed a significant association with age (β = 0.18 and 0.15; Bonferroni correction corrected p < 0.001), and parental education levels (β = 1.21 and 2.06; corrected p < 0.001). There were significant gender differences in fluid intelligence (male: 95.62 ± 17.31; female: 96.95 ± 17.16; β = 1.48, corrected p < 0.001), but not in crystallized intelligence (male: 107.38 ± 18.39; female: 106.80 ± 18.46; β = −0.44, corrected p > 0.05). Fluid intelligence showed a significantly positive association with household income (β = 0.09; corrected p = 0.013), but not crystallized intelligence (β = 0.06; corrected p > 0.05).

**Table 1:**
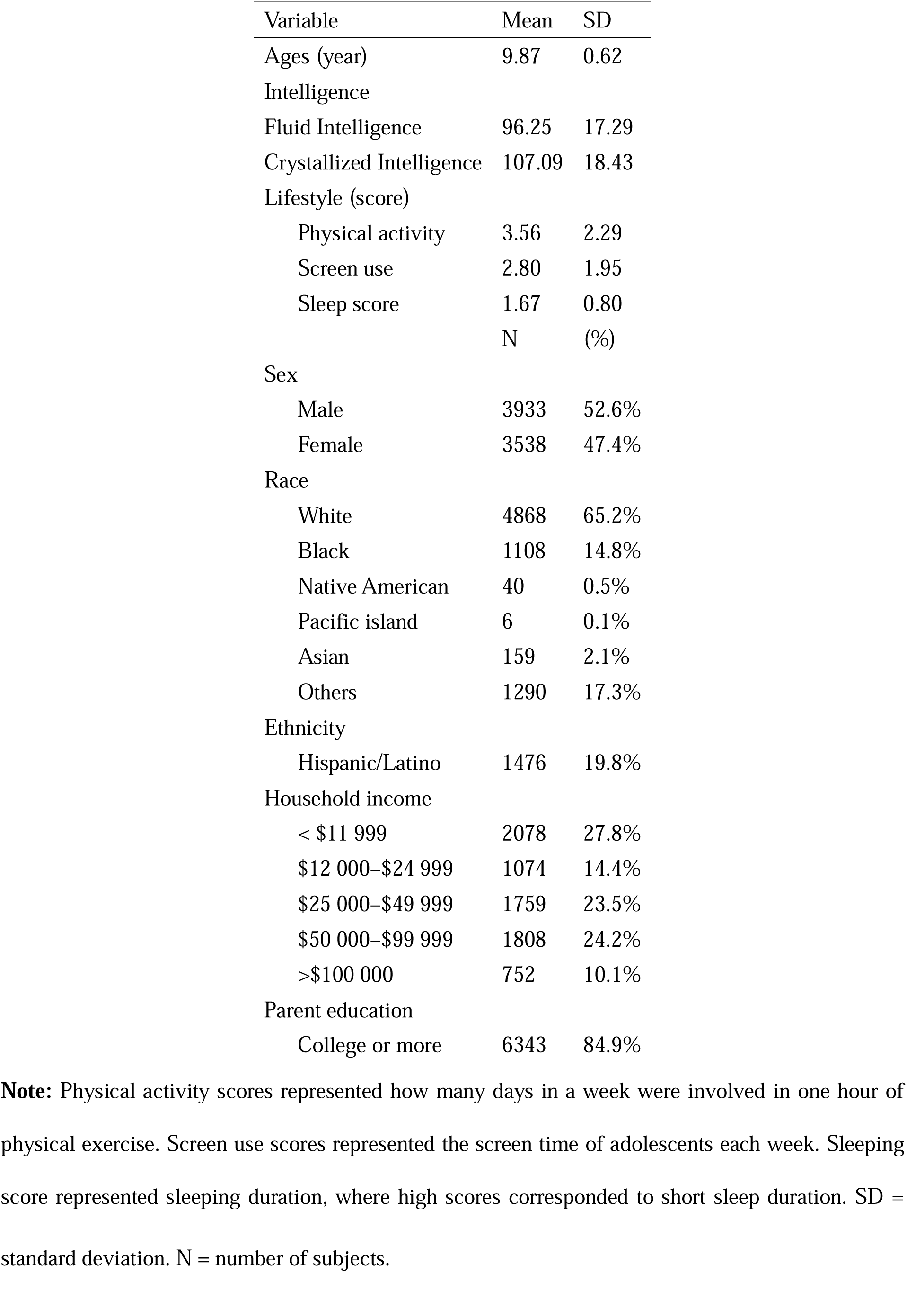
The demographic, intelligence and lifestyle statistics of the study population.

| Variable | Mean | SD |
| --- | --- | --- |
| Ages (year) | 9.87 | 0.62 |
| Intelligence |  |  |
| Fluid Intelligence | 96.25 | 17.29 |
| Crystallized Intelligence | 107.09 | 18.43 |
| Lifestyle (score) |  |  |
| Physical activity | 3.56 | 2.29 |
| Screen use | 2.80 | 1.95 |
| Sleep score | 1.67 | 0.80 |
|  | N | (%) |
| Sex |  |  |
| Male | 3933 | 52.6% |
| Female | 3538 | 47.4% |
| Race |  |  |
| White | 4868 | 65.2% |
| Black | 1108 | 14.8% |
| Native American | 40 | 0.5% |
| Pacific island | 6 | 0.1% |
| Asian | 159 | 2.1% |
| Others | 1290 | 17.3% |
| Ethnicity |  |  |
| Hispanic/Latino | 1476 | 19.8% |
| Household income |  |  |
| < \$11 999 | 2078 | 27.8% |
| \$12 000–\$24 999 | 1074 | 14.4% |
| \$25 000–\$49 999 | 1759 | 23.5% |
| \$50 000–\$99 999 | 1808 | 24.2% |
| >\$100 000 | 752 | 10.1% |
| Parent education |  |  |
| College or more | 6343 | 84.9% |

**Note:** Physical activity scores represented how many days in a week were involved in one hour of physical exercise. Screen use scores represented the screen time of adolescents each week. Sleeping score represented sleeping duration, where high scores corresponded to short sleep duration. SD =
standard deviation. N = number of subjects.

**Table 2:**
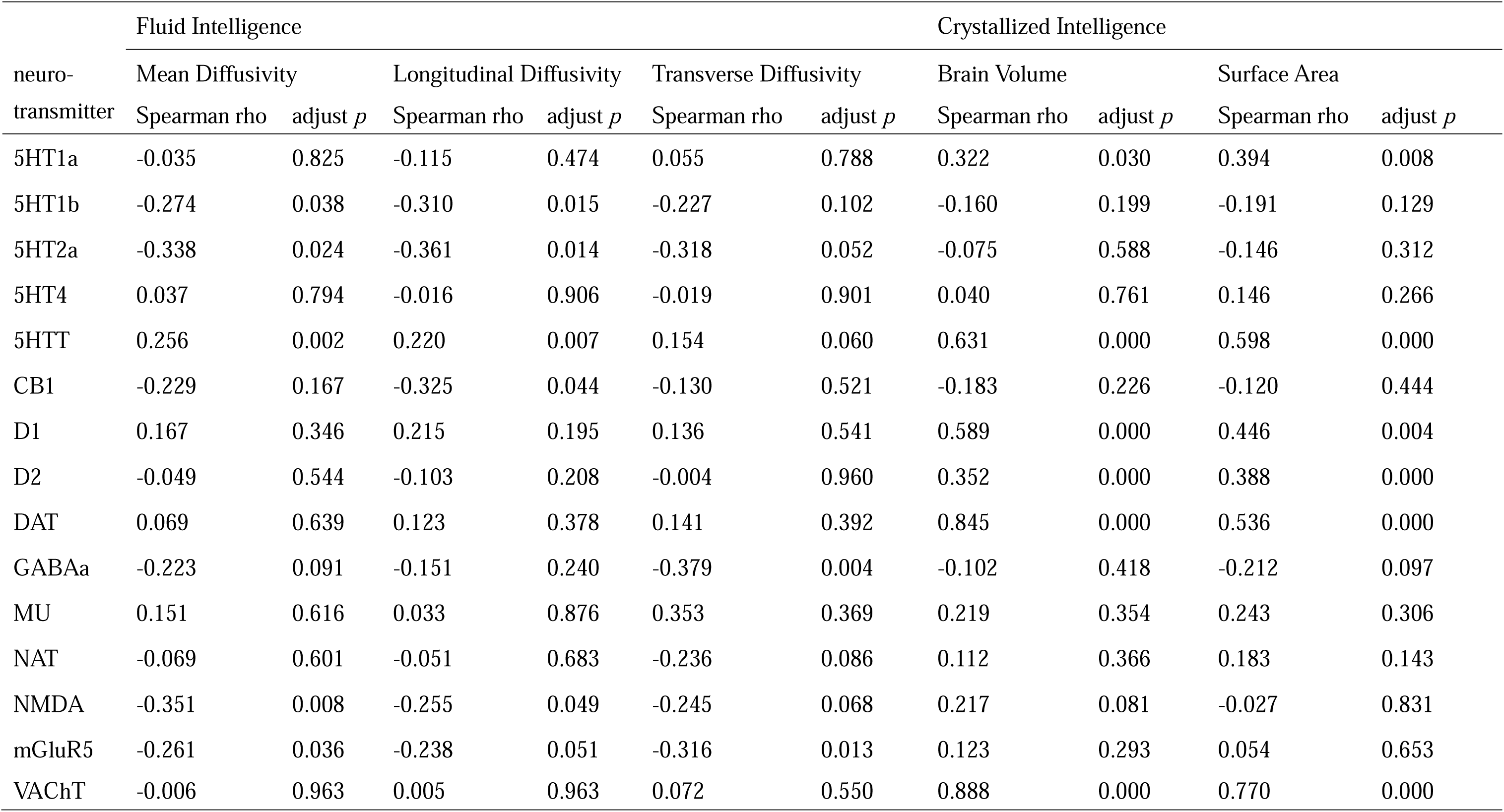
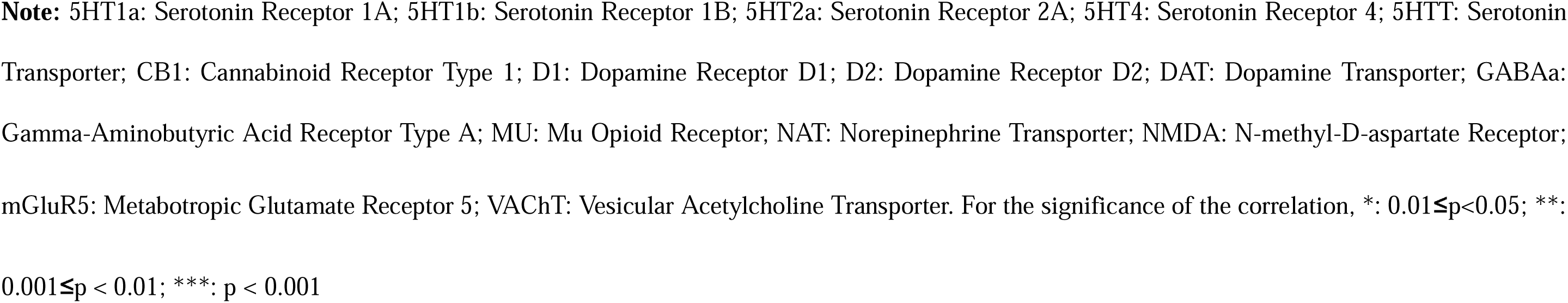
Neurotransmitters influencing fluid and crystallized Intelligence.

| neuro-transmitter | Fluid Intelligence |  |  |  |  |  | Crystallized Intelligence |  |  |  |
| --- | --- | --- | --- | --- | --- | --- | --- | --- | --- | --- |
|  | Mean Diffusivity |  | Longitudinal Diffusivity |  | Transverse Diffusivity |  | Brain Volume |  | Surface Area |  |
|  | Spearman rho | adjust *p* | Spearman rho | adjust *p* | Spearman rho | adjust *p* | Spearman rho | adjust *p* | Spearman rho | adjust *p* |
| 5HT1a | -0.035 | 0.825 | -0.115 | 0.474 | 0.055 | 0.788 | 0.322 | 0.030 | 0.394 | 0.008 |
| 5HT1b | -0.274 | 0.038 | -0.310 | 0.015 | -0.227 | 0.102 | -0.160 | 0.199 | -0.191 | 0.129 |
| 5HT2a | -0.338 | 0.024 | -0.361 | 0.014 | -0.318 | 0.052 | -0.075 | 0.588 | -0.146 | 0.312 |
| 5HT4 | 0.037 | 0.794 | -0.016 | 0.906 | -0.019 | 0.901 | 0.040 | 0.761 | 0.146 | 0.266 |
| 5HTT | 0.256 | 0.002 | 0.220 | 0.007 | 0.154 | 0.060 | 0.631 | 0.000 | 0.598 | 0.000 |
| CB1 | -0.229 | 0.167 | -0.325 | 0.044 | -0.130 | 0.521 | -0.183 | 0.226 | -0.120 | 0.444 |
| D1 | 0.167 | 0.346 | 0.215 | 0.195 | 0.136 | 0.541 | 0.589 | 0.000 | 0.446 | 0.004 |
| D2 | -0.049 | 0.544 | -0.103 | 0.208 | -0.004 | 0.960 | 0.352 | 0.000 | 0.388 | 0.000 |
| DAT | 0.069 | 0.639 | 0.123 | 0.378 | 0.141 | 0.392 | 0.845 | 0.000 | 0.536 | 0.000 |
| GABAa | -0.223 | 0.091 | -0.151 | 0.240 | -0.379 | 0.004 | -0.102 | 0.418 | -0.212 | 0.097 |
| MU | 0.151 | 0.616 | 0.033 | 0.876 | 0.353 | 0.369 | 0.219 | 0.354 | 0.243 | 0.306 |
| NAT | -0.069 | 0.601 | -0.051 | 0.683 | -0.236 | 0.086 | 0.112 | 0.366 | 0.183 | 0.143 |
| NMDA | -0.351 | 0.008 | -0.255 | 0.049 | -0.245 | 0.068 | 0.217 | 0.081 | -0.027 | 0.831 |
| mGluR5 | -0.261 | 0.036 | -0.238 | 0.051 | -0.316 | 0.013 | 0.123 | 0.293 | 0.054 | 0.653 |
| VACHT | -0.006 | 0.963 | 0.005 | 0.963 | 0.072 | 0.550 | 0.888 | 0.000 | 0.770 | 0.000 |

**Note:** 5HT1a: Serotonin Receptor 1A; 5HT1b: Serotonin Receptor 1B; 5HT2a: Serotonin Receptor 2A; 5HT4: Serotonin Receptor 4; 5HTT: Serotonin Transporter; CB1: Cannabinoid Receptor Type 1; D1: Dopamine Receptor D1; D2: Dopamine Receptor D2; DAT: Dopamine Transporter; GABA<sub>A</sub>: Gamma-Aminobutyric Acid Receptor Type A; MU: Mu Opioid Receptor; NAT: Norepinephrine Transporter; NMDA: N-methyl-D-aspartate Receptor; mGluR5: Metabotropic Glutamate Receptor 5; VACHT: Vesicular Acetylcholine Transporter. For the significance of the correlation, \*: $0.01 \leq p < 0.05$ ; \*\*: $0.001 \leq p < 0.01$ ; \*\*\*: $p < 0.001$

### MRI signatures of fluid and crystallized intelligence

The correlation between the fluid/crystallized intelligence and neuroimaging phenotypes including sMRI imaging data and dMRI imaging data using the LME model. The results of the LME analysis were shown in Supplement Table S1∼S7 for the two intelligence measures separately. In the separate LME model, we found that fluid and crystallized intelligence were significantly associated with cortical SA (fluid: 0.001 < β < 0.019, N = 66 of 148 ROIs; crystallized: 0.003 < β < 0.020, N = 134; Bonferroni correction corrected p < 0.05). Meanwhile, the significant association between fluid/crystallized intelligence and MD was distributed in similar regions. The overlap between these associations might be attributed to a significant correlation between the two forms of intelligence (r = 0.396, p < 0.001).

After controlling the influence of the crystallized intelligence in a combined LME model, MD of 24 cortical regions (-50.81 < β < -30.64, Bonferroni correction corrected p < 0.05; mainly in supramarginal gyrus, temporal gyrus, lateral sulcus, central sulcus, and occipital sulcus; Figure 2b) were significantly associated with fluid intelligence. LD of 8 cortical regions (-39.14 < β < -25.60, corrected p < 0.05; Figure S1f) and TD of 31 cortical regions (-56.89 < β < -27.10, corrected p < 0.05; Figure S1 g) were significantly associated with fluid intelligence. FA showed significant association only with left accumbens area (β = -13.20, corrected p < 0.05; Figure S1d). There were almost no significant associations between fluid intelligence and sMRI phenotypes except for one cortical region (right superior occipital sulcus and transverse occipital sulcus: β = 4.49×10^-3^, corrected p < 0.05; Figure 2a), BV of one cortical region (posterior corpus callosum: β = 4.62×10^-3^, corrected p < 0.05; Figure S1a), and CT of two cortical regions (left anterior part of the cingulate gyrus and sulcus: β = -5.92×10^-3^, corrected p < 0.05; left suborbital sulcus: β = -4.46×10^-3^, corrected p < 0.05; Figure S1c).

**Figure 2:**
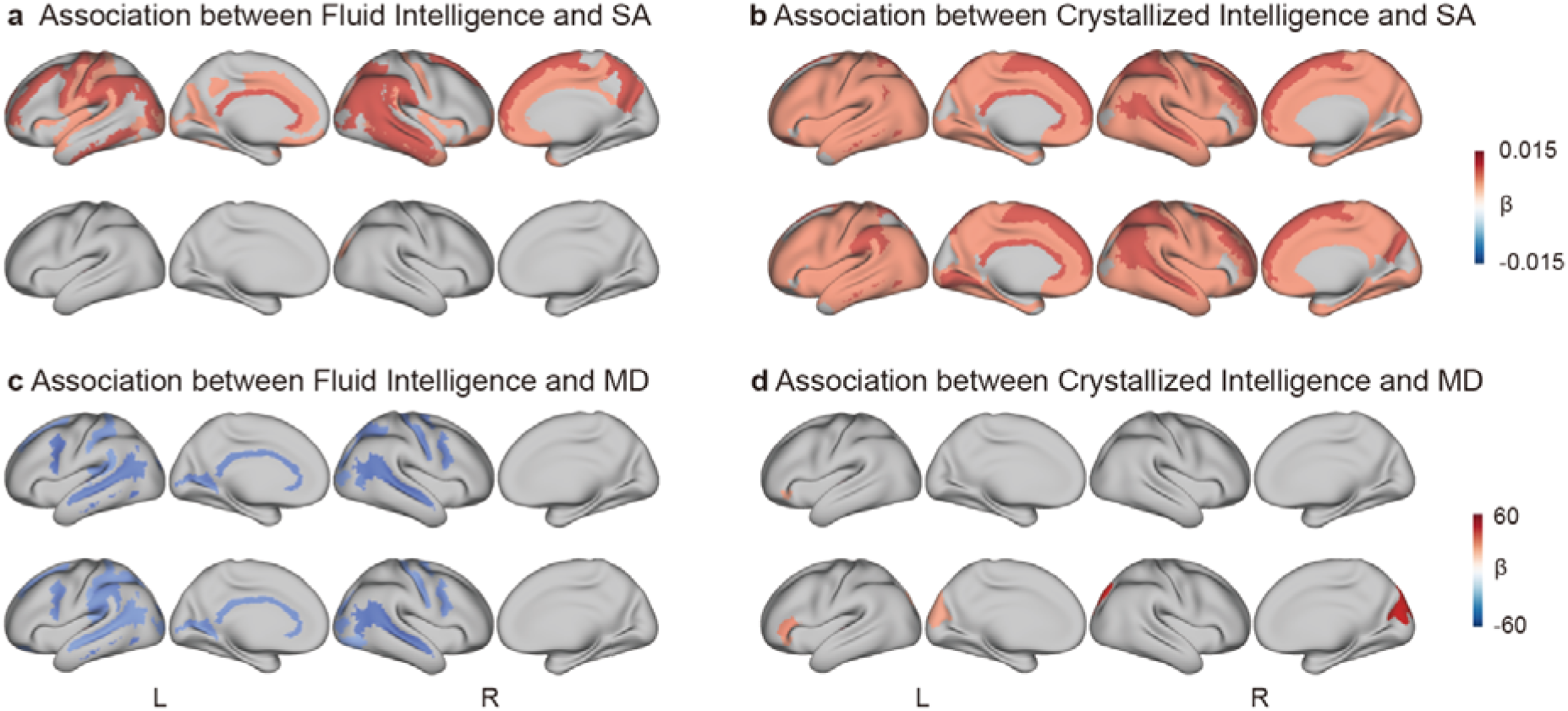
The associations between fluid/crystallized intelligence and neuroimaging phenotypes. The colored brain areas represented the significant associations after Bonferroni correction, and the value indicated the β-value between intelligence measures and MRI markers. In each panel, the upper row was the result of a separate LME model which retained the influence of the other form of intelligence; and the bottom row was the result of the combined LME model which removed the influence of the other form of intelligence. The associations between fluid/crystallized intelligence and other brain phenotypes are shown in Supplement Figure S1.

On the contrary, crystallized intelligence was significantly associated with sMRI phenotypes. After controlling the influence of fluid intelligence using LME model 2, BV of 127 cortical regions (6.27×10^-4^ < β < 8.65×10^-3^, corrected p < 0.05; Figure S1a) and SA of 126 cortical regions (0.0020 < β < 0.0244, corrected p < 0.05; Figure 2a) across the whole brain cortical gray matter were associated with crystallized intelligence. CT showed a significant association in one cortical region (left lingual gyrus: β = 1.34, corrected p < 0.05; Figure S1c). A small number of dMRI phenotypes were found to be associated with crystallized intelligence, including MD of 6 cortical regions (29.43 < β < 39.15, corrected p < 0.05; mainly in the inferior frontal gyrus, middle occipital gyrus, and cuneus; Figure 2b) and TD of 18 cortical regions (22.86 < β < 61.20, corrected p < 0.05; Figure S1g).

### Genetic Basis of Fluid and Crystallized Intelligence

Utilizing the association between MD and fluid intelligence as well as the association between SA and crystallized intelligence, the MRI-intelligence association maps (Figures 2c and b) were spatially correlated with the gene expression maps, followed by GO and KEGG enrichment analysis (Supplement Table S8∼S19). Through GO enrichment (Figures 3a and c), both fluid and crystallized intelligence demonstrated involvement in ion channels and signaling pathways that are crucial for neural communication and information processing. These genes participated in various ion channel activities (1.70 < fold enrichment < 7.52; q < 0.05), facilitating the passage of ions across cellular membranes essential for neuronal excitability and synaptic transmission. Additionally, they were associated with signaling pathways regulating synaptic function and neurotransmitter release (1.88 < fold enrichment < 2.98; q < 0.05), contributing to cognitive processes such as learning and memory. Processes such as amide transport (fold enrichment = 2.21 and 2.11; q < 0.05) and organic acid binding (fold enrichment = 2.05 and 2.02; q < 0.05) underscored their roles in cellular metabolism and neurotransmitter function, while involvement in synapse organization (fold enrichment = 2.19 and 1.88; q < 0.05) and cell projection membrane (fold enrichment = 1.91 and 2.10; q < 0.05) highlighted their contribution to intercellular communication and synaptic connectivity.

**Figure 3:**
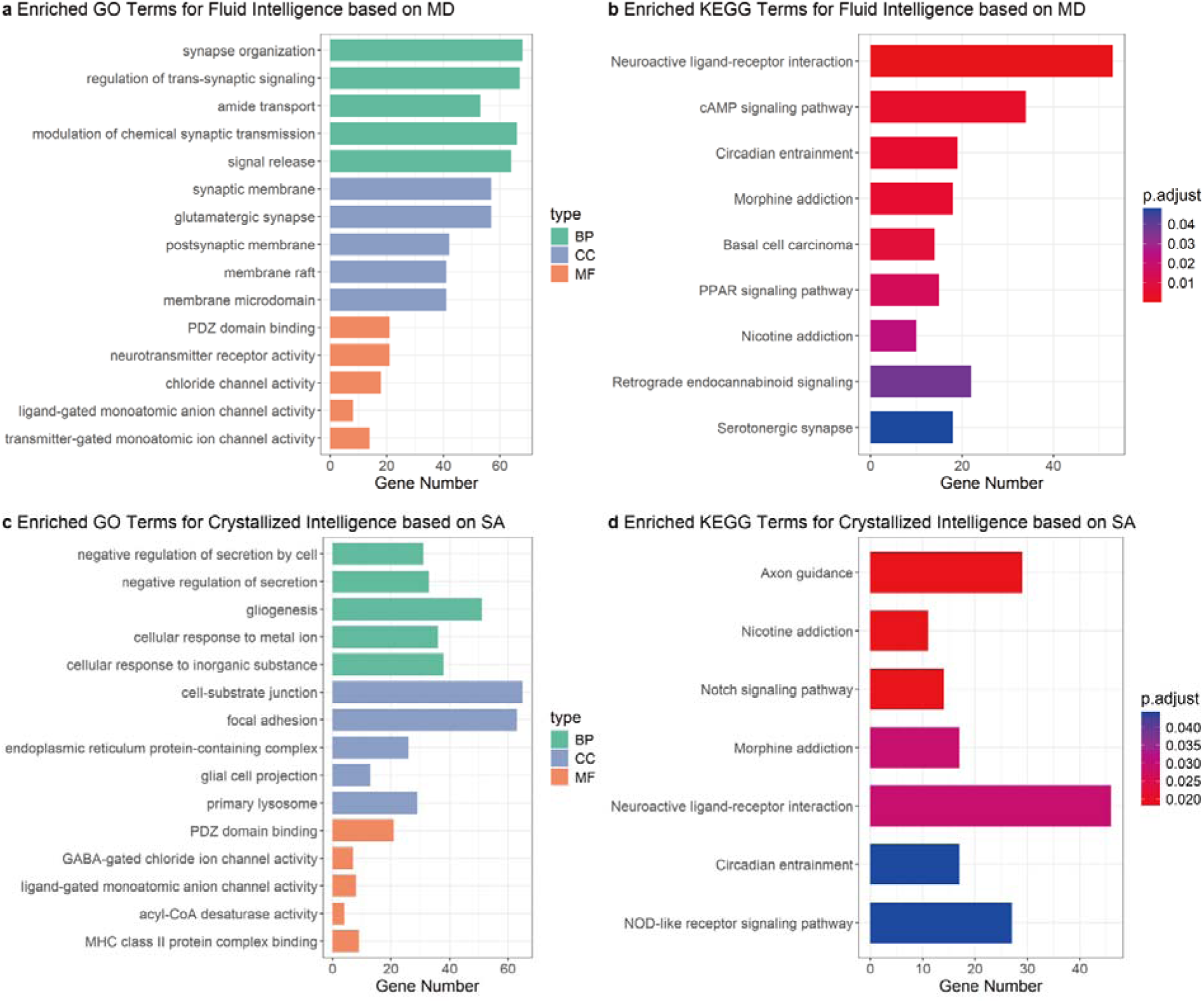
The results of PLRS and GO/KEGG enrichment of fluid and crystallized intelligence. (a) (b)The results of the GO/KEGG enrichment of fluid-intelligence-related genes based on MD (c)(d)The results of the GO/KEGG enrichment of crystallized-intelligence-related genes based on SA. BP = Biological Process, CC = Cellular Component, MF = Molecular Function

**Figure 4:**
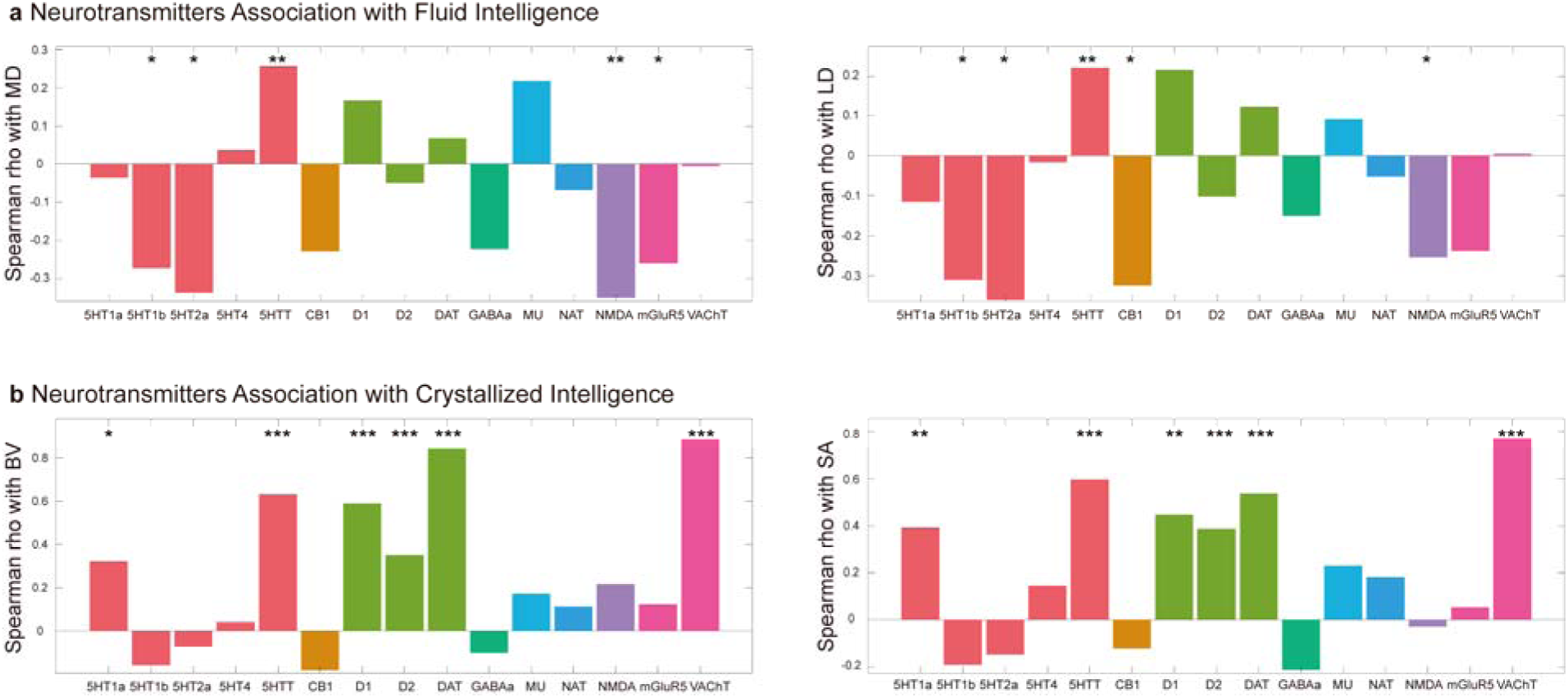
Neurotransmitters influencing fluid and crystallized Intelligence. (a) Spearman rho of the spatial correlation between fluid-intelligence-MD association map and the neurotransmitter density maps. (b) Spearman rho of the spatial correlation between crystallized-intelligence-BV association map and the neurotransmitter density maps. *: 0.01≤p<0.05; **: 0.001≤p < 0.01; ***: p<0.001

The complexity of synaptic and membrane-related functions varied between fluid and crystallized intelligence. Fluid intelligence exhibited a more complicated relationship with these processes, including specific functions such as glutamatergic synapse function, postsynaptic membrane activity, and membrane raft organization (fold enrichment = 2.10, 2.29, and 2.15; q < 0.05). This complexity may be explained by the dynamic and adaptive nature of fluid intelligence, characterized by rapid information processing and cognitive flexibility. In contrast, crystallized intelligence demonstrated a more straightforward involvement in synaptic and membrane-related functions, suggesting a more stable and structured cognitive processing. Furthermore, crystallized intelligence is uniquely associated with protein-related functions, such as endoplasmic reticulum protein-containing complex, MHC class II protein complex binding, and scaffold protein binding (fold enrichment = 3.05, 4.66 and 2.88; q < 0.05), implicating roles in protein synthesis, immune response modulation, and structural scaffolding supporting crystallized intelligence-specific cognitive processes.

Through KEGG enrichment (Figure 3b, d), both fluid and crystallized intelligence exhibited involvement in environmental information processing, particularly in neuroactive ligand-receptor interaction (fold enrichment = 1.83 and 1.69; q < 0.05), and shared associations with addictive behaviors, indicated by relevance to Morphine addiction (fold enrichment = 2.75 and 2.46; q < 0.05) and Nicotine addiction (fold enrichment = 3.47 and 3.62; q < 0.05). Distinctions emerged in the organismal systems category, where fluid intelligence was associated with the cAMP and PPAR signaling pathways (fold enrichment = 2.04 and 2.78; q < 0.05), while crystallized intelligence was linked to the Notch and NOD-like receptor signaling pathways (fold enrichment = 2.98 and 1.91; q < 0.05), indicating divergent pathways related to intracellular signaling and immune response modulation. Additionally, fluid intelligence demonstrated relevance to the GABAergic synapse (fold enrichment = 2.34; q < 0.05), suggesting relevance to neurotransmission, while crystallized exhibited associations with axon guidance (fold enrichment = 2.10; q < 0.05), suggesting relevance in neuronal development.

Enrichment analyses of LD- and TD-related genes showed similar results to those using MD in GO and KEGG terms. Similarly, analyses based on BV and SA also showed comparable results in these terms. Details could be found in the Supplement Table S8∼S19.

### Neurotransmitter Basis of Fluid and Crystallized Intelligence

Spatial correlation between intelligence-associated MRI feature maps and neurotransmitter density maps was performed to obtain the neurotransmitter basis of two forms of intelligence. For fluid intelligence, we mainly focused on MD and LD and found their association maps were significantly associated with the neurotransmitter density of the serotonin system, glutamate system, and GABA system. The associated serotonin neurotransmitters included 5HT1b (Spearman rho = −0.310, −0.274; adjusted p = 0.015, 0.038), 5HT2a (Spearman rho = −0.361, −0.388; p = 0.014, 0.024) and 5HTT (Spearman rho = 0.220, 0.256; p = 0.007, 0.002). The associated glutamate neurotransmitters included mGluR5 (Spearman rho = −0.238, −0.261; p = 0.051, p = 0.036) and NMDA (Spearman rho = −0.255, −0.351; p = 0.049, 0.008). Additionally, based on TD, fluid intelligence was significantly associated with GABAa (z = −0.379; p = 0.005).

For crystallized intelligence, we mainly assessed crystallized-intelligence-BV/SA association maps and revealed their significant associations with serotonin, dopamine, and acetylcholine systems. The associated serotonin neurotransmitters included 5HT1a (Spearman rho = 0.322, 0.394; adjusted p = 0.030, 0.008) and 5HTT (Spearman rho = 0.631, 0.598; both p < 0.001). The associated dopamine neurotransmitters included D1 (Spearman rho = 0.589, 0.446; p < 0.001, p = 0.004), D2 (Spearman rho = 0.352, 0.388; both p < 0.001) and DAT (Spearman rho = 0.845, 0.536; p < 0.001). The associated acetylcholine neurotransmitter was VAChT (Spearman rho = 0.888, 0.770; both p < 0.001).

### Impact of lifestyle on two forms of intelligence and the neural mediates

According to the 24-Hour Movement Guidelines, we included physical activities, screen use, and sleep duration in the analysis. Physical activity days per week showed significant associations with fluid and crystallized intelligence (β = 0.30, p = 7.43×10^−4^; β = 0.18, p = 0.042). Using mediation analysis, SA of 32 cortical regions (0.011 < AME < 0.035, p < 0.05; Figure 5a and Supplement Table S20) exhibited significant full mediating effects on the association between physical activities and crystallized intelligence. The spatial pattern of these significant mediating effects in the brain (primarily distributed in the temporal, frontal, and parieto-occipital cortex) was associated with the distribution of the serotonin system (5HT1a: Spearman rho = 0.340, adjusted p = 0.005) and dopamine system (D2: Spearman rho = 0.187, p = 0.022).

**Figure 5:**
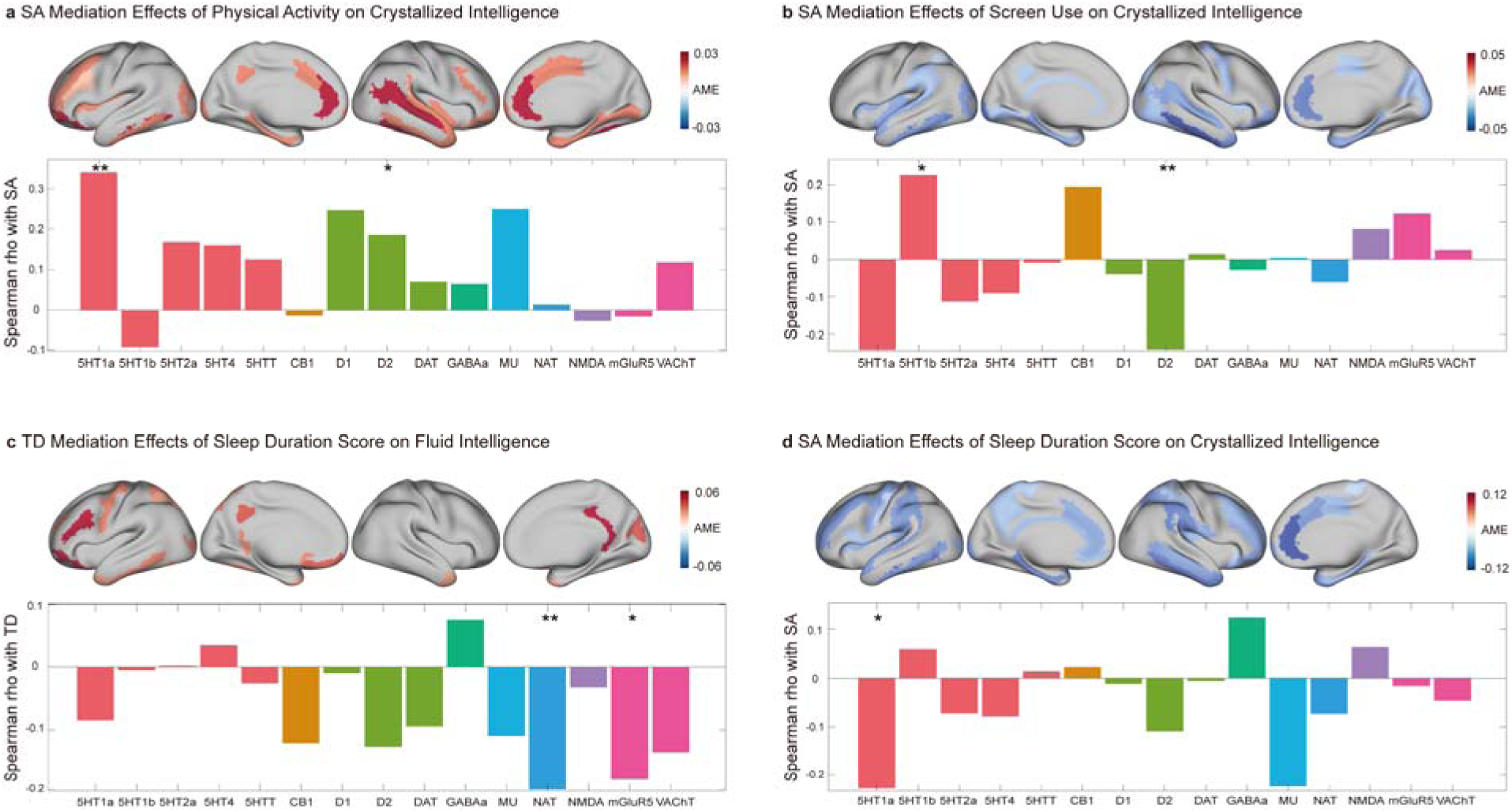
The mediation effect of lifestyle and neural basis of the mediation. In each panel, the upper row was the average mediation effect of the mediation model; the bottom row was a result of the spatial correlation between mediation maps and neurotransmitter maps. *: 0.01≤p<0.05; **: 0.001≤p < 0.01; ***: p<0.001

Screen use showed negative associations with fluid and crystallized intelligence (β = –0.30, p = 0.005; β = –0.53, p = 6.4×10^−7^). SA of 43 cortical regions (-0.05 < AME < -0.014, p < 0.05; Figure 5b and Supplement Table S21) exhibited significant mediating effects on the association between screen time and crystallized intelligence. The spatial pattern of these mediating effects (mainly distributed in the temporal, occipital, and parieto-occipital cortex) was significantly associated with serotonin system (5HT1b: Spearman rho = 0.225, adjusted p = 0.029) and dopamine system (D2: Spearman rho = –0.240, p = 0.004).

Sleep duration score (β = –0.56, p = 0.038; β = –1.32; p = 1.21×10^−6^) had negative associations with both fluid and crystallized intelligence. TD of 22 cortical regions (0.03 < AME < 0.06, p < 0.05; Figure 5c and Supplement Table S22) exhibited significant mediating effects on the association between short sleep duration and fluid intelligence. On the other hand, SA of 52 cortical regions (-0.12 < AME < -0.03, p < 0.05; Figure 5d and Supplement Table S23) exhibited significant mediating effects on the association between short sleep duration and crystallized intelligence. The spatial pattern of the mediating effects on fluid intelligence was significantly associated with norepinephrine system (NAT: Spearman rho = –0.201; p = 0.009) and glutamate system (mGluR5: Spearman rho = –0.182; p = 0.025). Conversely, the mediating pattern relating to crystallized intelligence was significantly associated with serotonin system (5HT1a: Spearman rho = –0.233, adjusted p = 0.046). These neurotransmitters with mediation effects agreed well with aforementioned findings.

## Discussion

In this study, imaging and intelligence data from the ABCD study allowed us to understand the specific neural signature and molecular basis of fluid and crystallized intelligence. We found that fluid intelligence was significantly associated with cortical microstructural phenotypes such as MD in the supramarginal gyrus, temporal gyrus, lateral sulcus, central sulcus, and occipital sulcus. Crystallized intelligence was significantly associated with macrostructural phenotypes such as SA in almost all cortical regions. Through spatial correlation with AHBA and neurotransmitter maps, we found that fluid and crystallized intelligence had their distinct neural basis while also sharing overlapping components. Fluid intelligence had a significant association with the glutamate system (receptor mGluR5 and NMDA) and GABA system (receptor GABAa), whereas crystallized intelligence had a significant association with the dopamine system (receptor D1, D2, and transporter DAT) and acetylcholine system (transporter VAChT). Through mediation analysis, we found that the way by which sleep affects fluid intelligence was mediated by the glutamate system, and the mediation effects of exercise, screen use, and sleep on crystallized intelligence were related to the dopamine and serotonin systems. These findings imply that keeping a healthy lifestyle may help to enhance the development of intelligence in adolescents.

Previous studies have revealed that fluid intelligence is linked to the brain volume and surface area of the dorsolateral prefrontal cortex, anterior cingulate gyrus, and inferior and superior parietal lobules ^8,10,12^. Meanwhile, higher crystallized intelligence level was found associated with higher whole-brain gray matter volume and cortex thickness ^10,13–15^. In this study, we applied two LME models that retained or removed the influence of the other form of intelligence. The separate LME model closely resembled previous approaches to the Human Connectome Project ^73^, and reached similar results that MRI phenotypes of fluid and crystallized intelligence were highly overlapped ^74^. The combined LME model indicated no significant associations between fluid intelligence and sMRI phenotypes. Compared to the separate LME model, we speculated that the association between the aforementioned MRI markers and fluid intelligence was largely due to the correlation between fluid and crystallized intelligence.

In our study, significant associations between fluid intelligence and cortical microstructural phenotypes were observed specifically within the supramarginal gyrus, temporal gyrus, lateral sulcus, central sulcus, and occipital sulcus, aligning with the Parieto-Frontal Integration Theory (P-FIT) which emphasized the importance of neural integration across parietal and frontal areas ^12^. The supramarginal gyrus supported phonological processing and working memory ^75,76^, essential for complex problem-solving integral to fluid intelligence. The temporal gyrus, involved in language comprehension ^77,78^, underpinned abstract thinking and memory integration, crucial for applying new knowledge. The lateral and central sulci, associated with sensory and motor processing ^79^, might facilitate the coordination required for executing complex operations. Similarly, the occipital sulcus contributed to visual-spatial reasoning ^80^, vital for manipulating spatial information. Together, these findings suggested that fluid intelligence was underpinned by a network of cortical areas, each contributing to the overall cognitive flexibility and problem-solving capabilities, reflected in the microstructural integrity of these regions. The result supported the viewpoint that fluid intelligence involved brain functions linked through an efficient parietal-frontal network.

For crystallized intelligence, the positive associations between crystallized intelligence and whole-brain cortical macro-structures were aligned with the Neural Efficiency Hypothesis ^81^ and concepts of brain and cognitive reserve ^82,83^. Higher intelligence correlated with more efficient neural processing, allowing for quicker access and use of learned skills and knowledge. Well-developed cortical architecture, greater cortical mass, and integrity were better equipped for efficient neural transmission and processing speed in cognitive challenges. Furthermore, we found that fluid intelligence was closely associated with the glutamate system (receptor mGluR5 and NMDA), GABA system (receptor GABAa), and serotonin system (transporter 5HTT, receptors 5HT1b and 5HT2a). The findings aligned with previous studies where a higher working memory index was significantly correlated with a lower GABA level ^30,31,84^ and lower GABA to glutamate ratio levels ^28^, as well as other studies which reported cognitive function was related to serotonin system ^20^ in transporter ^21,22^ and receptors ^85,86^. These significant associations suggested complex mechanisms where the capacity for fluid intelligence was influenced by both neural excitation and inhibition provided by glutamate and GABA, as well as the modulatory effects of serotonin on cognitive processes.

Crystallized intelligence had a significant association with the dopamine system, acetylcholine system, and serotonin system. In the formation of crystallized intelligence, acetylcholine enhanced the encoding of new information into long-term memory ^87^, establishing a base for memory formation. Dopamine helped quickly pull up stored knowledge when necessary ^88^, aiding in decision-making and problem-solving tasks. Meanwhile, serotonin enriched these memories with emotional and contextual details during the consolidation phase ^89^, making them more relatable and easier to recall. Together, these neurotransmitters form a sophisticated cognitive system that helps use learned knowledge in various situations.

Not only the neurotransmitters play a determinant role, but lifestyle in adolescents also influences the development of intelligence. Previous research has found that sleep and physical exercise are beneficial for intelligence ^38,39^, while screen time has adverse effects on intelligence. ^41–45^. Our findings were consistent with these findings. The mGlu5 activity was important for synaptic adjustments and cognitive processes associated with fluid intelligence ^90^. This suggested that sleep might facilitate these processes by optimizing receptor mGlu5 functionality, and thus enhancing fluid intelligence by improving synaptic connections and neural communication critical for learning and problem-solving tasks. The receptor 5HT1a in the Serotonin system impacted cognitive areas like learning, memory retention, and knowledge recall ^91^, which were key to crystallized intelligence. Serotonin levels during sleep affected the cognitive processing of learned knowledge, and thus effective serotonin regulation during sleep via receptor 5HT1a could be vital for utilizing long-term memory and knowledge for crystallized intelligence enhancement. The interaction between Dopamine receptor D2 and both physical activity and screen use showed different effects on crystallized intelligence. Physical activity positively stimulated receptor D2, enhancing motor flexibility and the ability to learn motor skills. This could improve abilities related to physical tasks and movements ^92^. On the other hand, too much screen time could overstimulate receptor D2, potentially harming crystallized intelligence. This over-stimulation might reduce attention and information processing efficiency, making it harder to use learned knowledge and skills effectively. The receptor D2 played a complex role in shaping crystallized intelligence, enhancing it through physical activity and potentially impairing it with excessive screen use.

## Conclusion

This study identified distinct neuroimaging, genetic, and neurotransmitter bases between fluid and crystallized intelligence. Our findings highlighted microstructural phenotypes for fluid intelligence and macrostructural phenotypes for crystallized intelligence as respective biomarkers. Molecular mechanisms derived from the aforementioned biomarkers suggested that fluid intelligence mainly had a significant association with the serotonin and glutamate system; while crystallized intelligence was related to serotonin, dopamine, and acetylcholine systems. Lifestyles including physical activities had positive associations but screen use and short sleep duration had negative associations with fluid and crystallized intelligence, mediated by mGlu5 receptors for fluid intelligence and 5HT1a and D2 receptors for crystallized intelligence. These findings provided insights into the neurobiological underpinnings of fluid and crystallized intelligence, laying the groundwork for subsequent observational or interventional research on intelligence.

