## Supplement Figure for "Neural correlates differ between crystallized and fluid intelligence in adolescents"

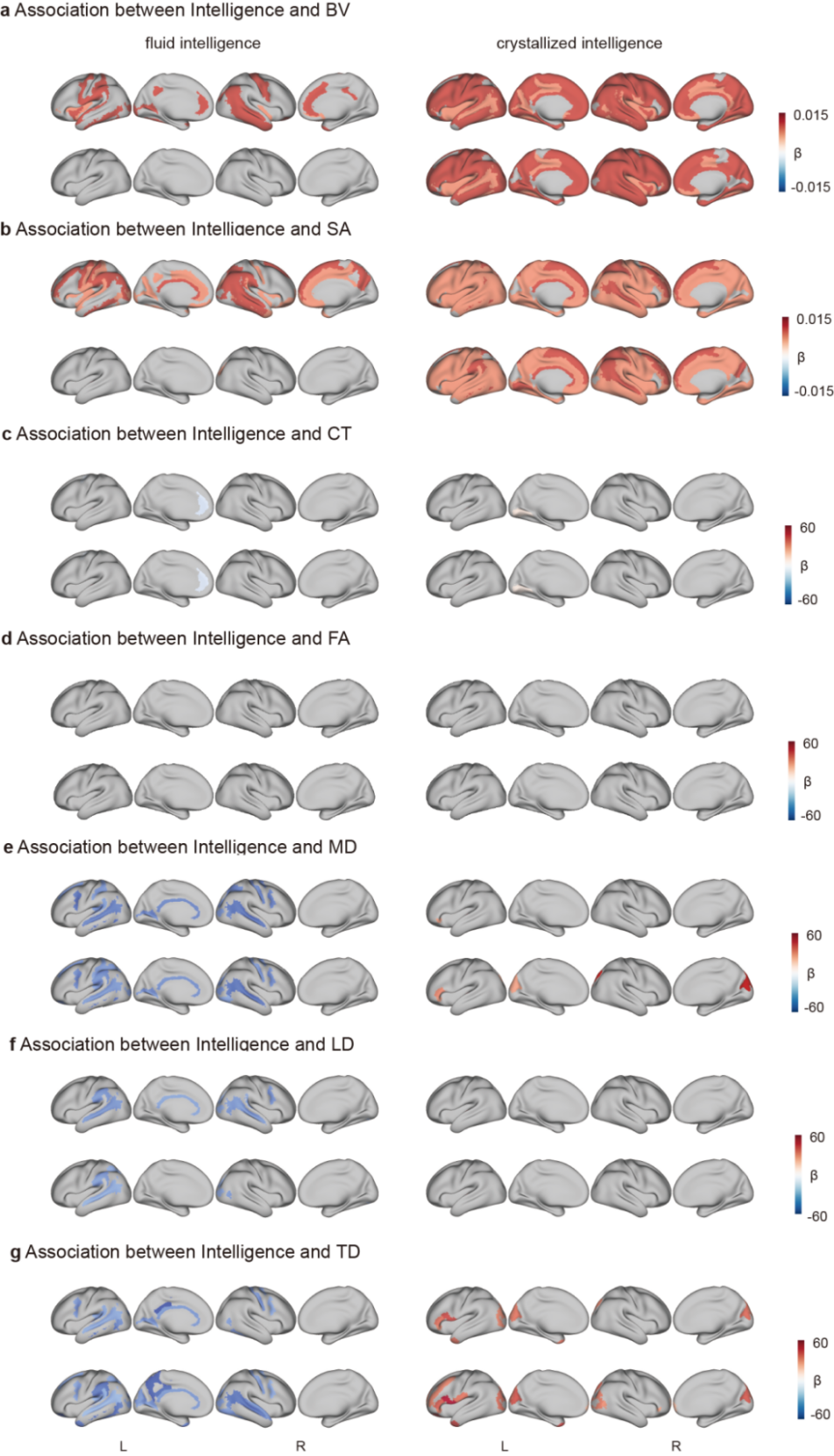

**Supplement Figure S1:** The associations between fluid/crystallized intelligence and MRI phenotypes using two types of LME models**.** The colored brain area represented the significant results after Bonferroni correction, and the value indicated the β-value between intelligence and MRI phenotype. In each panel the upper row was based on separate LME model, and the bottom row was based on combined LME model.
